# High-Voltage Biomolecular Sensing Using a Bacteriophage Portal Protein Covalently Immobilized Within a Solid-State Nanopore

**DOI:** 10.1101/2022.08.07.503088

**Authors:** Mehrnaz Mojtabavi, Sandra J. Greive, Alfred A. Antson, Meni Wanunu

## Abstract

The application of nanopores as label-free, single-molecule biosensors for electrical or optical probing of structural features in biomolecules has been widely explored. While biological nanopores (membrane proteins and bacteriophage portal proteins) and solid-state nanopores (thin films and two-dimensional materials) have been extensively employed, the third class of nanopores known as hybrid nanopores, where an artificial membrane substitutes the organic support membrane of proteins, has been only sparsely studied, due to challenges in implementation. *G20c* portal protein contains a natural DNA pore that is used by viruses for filling their capsid with viral genomic DNA. We have previously developed a lipid-free hybrid nanopore by “corking” the *G20c* portal protein into a SiN_*x*_ nanopore. Herein, we demonstrate that through chemical functionalization of the synthetic nanopore, covalent linkage between the solid-state pore and the *G20c* portal protein considerably improves the hybrid pore stability, lifetime, and voltage resilience. Moreover, we demonstrate electric-field-driven and motor protein-mediated transport of DNA molecules through this hybrid pore. Our integrated protein/solid-state hybrid nanopore can serve as a robust and durable framework for sensing and sequencing at high voltages, potentially providing higher resolution, higher signal-to-noise ratio, and higher throughput compared to the more conventional membrane-embedded protein platforms.

## Introduction

Studying biomolecular systems at the single-molecule level has unleashed a new era in probing the structure and dynamics of biomolecules with unprecedented detail. Nanopore sensors have shown unprecedented performance in recognition^1-9^ and quantification^10-16^ of various analytes and sequencing of individual molecules.^17-19^ Progress in nanopore technology in the last decade has resulted in the development of two platforms: solid-state nanopores^20-27^ and protein nanopores,^28-34^ with each platform having specific advantages in terms of performance/resolution, signal reproducibility, pore size/chemistry tunability, and chemical/physical robustness. Protein nanopores typically require an organic support membrane (e.g., lipid bilayers or block-copolymers) that is significantly less robust than ceramic-based solid-state membranes (e.g., silicon nitride). On the other hand, no methods exist for reproducing solid-state nanopores in atomic detail, preventing progress in solid-state nanopore technology. Combining the two types of pore systems to form hybrid pores^35, 36^ has been achieved, although both reports shared the same limitation, namely, the protein pore was not chemically linked to the solid-state pore, which results in a transient hybrid pore to which a limited voltage range can be applied. Site-specific, reproducible, and uniformly oriented protein immobilization within a synthetic membrane in a manner that preserves the protein conformation and functionality requires an understanding of the membrane’s surface chemistry, protein’s properties, and the nature of their interaction to eliminate non-specific binding. We recently hacked the *G20c* portal protein into insertion into a lipid bilayer environment (unnatural for the portal protein)^37^ and later demonstrated voltage-induced “corking” of the portal into a solid-state nanopore.^36^ This protein has shown high stability and tunability. In this work, we developed a novel type of hybrid nanopore system by covalently immobilizing the *G20c* portal protein within a chemically modified SiN_*x*_ nanopore to fabricate an ultra-stable hybrid nanopore that is functional at high voltages. We show the application of this hybrid nanopore in sensing DNA molecules through free translocation and motor protein-mediated ratcheting at high voltages. The ability to apply high voltage in a protein-based nanopore platform can allow control over enzyme-based DNA movement, long-range DNA scanning, and further can be used for measurements of the pore resolution at voltages that are beyond the capabilities of traditional membrane-based pores. As we show here, our approach to producing hybrid nanopores by chemically linking protein to the solid-state pore constitutes the first step towards high-performance, high throughput array-based pore devices.

## Results and Discussion

*G20c* portal protein from the thermostable bacteriophage *G20c* oligomerizes to form a dodecameric assembly with an internal channel containing 1.8 nm constriction, as shown in **Figure 1a**. In common with the wild-type protein, its CD/N mutant, where four aspartic acid residues (D) were substituted by asparagine (N) to alter the internal surface charge for enhanced sensing of negatively charged molecules,^36^ in addition to substitution of an externally facing leucine residue to cysteine for chemical labeling,^37^ also forms 12-mers (**Figure S1**). In contrast to the most successfully implemented proteins as nanopore sensors that assemble to form channels upon insertion into the lipid bilayer or synthetic polymer membranes, the hydrophilic *G20c* portal protein is water-soluble and fully assembles in an aqueous solution. So far, our efforts have resulted in two distinct methods to utilize this protein as a nanopore sensor: 1. insertion into the lipid bilayer through a hydrophobic conjugate^37^ and 2: corking into solid-state nanopore to form a lipid-free hybrid nanopore.^36^ However, none of these methods have been efficient enough to deliver high-yield, full-range sensing capability in various experimental conditions, particularly failing at a high applied voltage.

**Figure 1.**
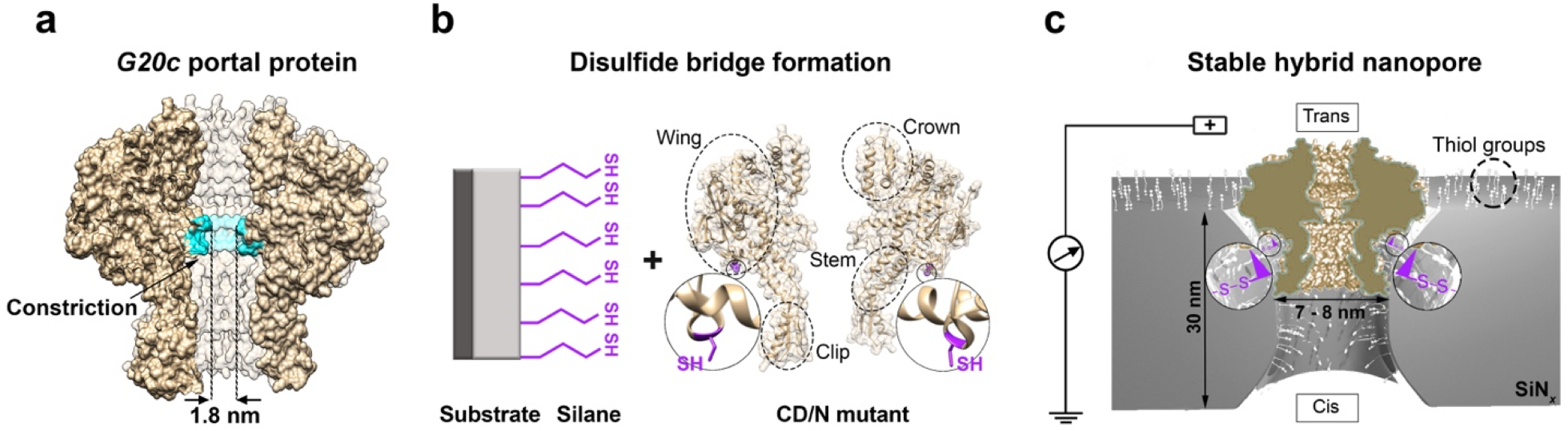
*G20c* portal protein immobilization within a chemically modified solid-state nanopore. (a) Molecular surface of the *G20c* portal protein showing 1.8 nm constriction formed by tunnel loops of 12 subunits. Only 6 subunits are shown to visualize the central tunnel (vertical). (b) Schematic for disulfide bridge formation between cysteine residues of the CD/N *G20c* portal protein and the thiolated substrate. For clarity only 2 subunits of the portal protein are shown. (c) Insertion of the CD/N *G20c* portal protein into the thiolated SiN*x* nanopore and formation of disulfide bridges between the protein and the pore wall ensures a highly stable hybrid nanopore.

Building upon our previous works, in this work we exploit the cysteine residues residing under the wing and near the cap of the protein to form disulfide bridges with a thiolated solid-state nanopore, as shown in **Figure 1b**. The ideal positioning of the cysteine mutation allows site-specific, reproducible, and uniformly oriented protein immobilization within the thiolated SiN_*x*_ nanopore while preserving its conformation and activity. We predict that the dominant protein orientation within the SiN_*x*_ nanopore, shown in **Figure 1c**, is required for the protein to cross-link to the chemically-modified substrate. Given this specific geometric point of the cysteine modification in the protein, formation of an undesirable disulfide bridge between the protein and the planar surface of the SiN_*x*_ membrane is not feasible.

In order to chemically modify the SiN_*x*_ membrane, we used silane coupling agents that form durable self-assembled monolayers (SAMs) through covalent bonding to SiO_x_-rich surfaces. SiN_*x*_ is also amenable to silanization because it contains a silicon-rich, nitrogen-depleted area on its surface which is terminated by oxide and hydroxide groups.^38^ This native oxide layer is beneficial for direct chemical modification of SiN_*x*_ nanopores with silanes.^39-41^ For our purpose, thiol functionality is imparted to the surface by treating the SiN_*x*_ membrane with 2,2-dimethoxy-1-thia-2-silacyclopentane (thia-silane). Thia-silane reacts rapidly with the surface through ring-opening click reaction driven by the difference in bond energies and relief of the ring strain. This reaction is simple, fast, high-yield, and occurs via cleavage of Si-S bond by hydroxyl groups as previously reported.^42-44^ The ring structure of the thia-silane protects the sulfhydryl group from forming disulfide bonds prior to SAM formation (**Figure 2a)**. In contrast, in other thiol-functionalized silanes such as 3-(mercaptopropyl)trimethoxysilane (MPTES), deposition on the substrate occurs through several steps of hydrolysis and condensation reactions that are more prone to form multi-layers on the surface (**Figure S2**). To chemically modify the SiN_*x*_ nanopores, 7-8 nm diameter pores were fabricated through a 30-nm thick SiN_*x*_ membrane with a focused transmission electron microscope (TEM) beam and thoroughly cleaned with piranha solution for any organic residue and contamination to be removed and the surface to be rendered hydrophilic and hydroxylated (see Methods Section). Afterward, the SiN_*x*_ membranes were incubated with silane solution for 1 - 3 hours. Silanized pores were rinsed with the solvent in several steps and assembled in a fluidic cell with two 70 µl volume chambers across the chip for trans-pore ion current measurements. Ion transport through a nanopore could be related to the pore geometry using a pore conductance model that takes into account pore dimensions and an access resistance term.^6^ Thia-silane molecules with an extended length of ∼ 7 Å reduce the pore diameter and increase its overall thickness by approximately 1.4 nm. Therefore, silanized pores are expected to have less conductance than expected from their TEM-measured diameter (d_TEM_), as shown in **Figure 2b**.

**Figure 2.**
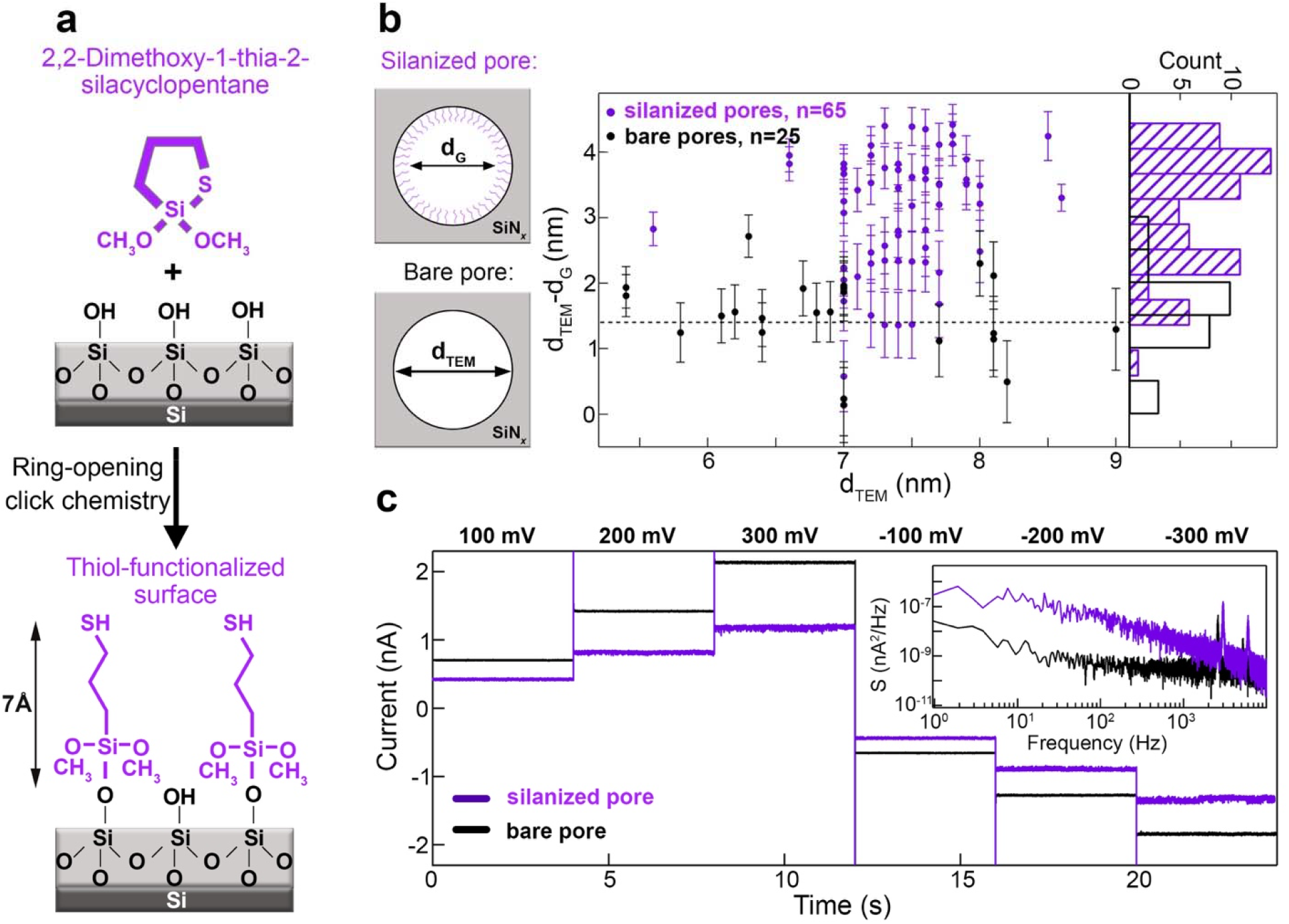
SiN_*x*_ nanopore chemical modification with cyclic thia-silane. (a) Functionalization of SiN_*x*_ surface with thia-silane through ring-opening click chemistry. (b) Nanopore diameter change after silanization results in decreased ionic conductance. d^’^_G_: nanopore diameter calculated from the nanopore conductance using the nanopore resistance model. d_TEM_: nanopore diameter from TEM image taken after fabrication. The plot shows d_TEM_ - d^’^_G_ vs. d_TEM_ demonstrating the deviation of d^’^_G_ from d_TEM_ for both bare and silanized SiN_*x*_ pores. The error bars show the values calculated with effective nanopore thickness ranging from 10 - 15 nm. (c) Ionic current trace of a 7 nm bare SiN_*x*_ nanopore shown in black and a 7 nm silanized SiN_*x*_ nanopore shown in purple. The inset shows PSD of both pores at 100 mV. Traces were recorded at 250 kHz sampling frequency and lowpass filtered at 10 kHz. Buffer: 0.5 M NaCl, 20 mM HEPES, pH 7.5.

We used the pore resistance model to calculate the bare and silanized SiN_*x*_ pore diameters from the ionic conductance after single-channel measurement (d^’^_G_) and compare it to their d_TEM_. **Figure 2b** shows a systematic deviation of d^’^_G_ from d_TEM_ for both bare and silanized SiN_*x*_ pores. While for the bare SiN_*x*_ pores, the deviation is 1.51 ± 0.63 nm, for silanized pores we find a deviation of 3.1 ± 0.9 nm. While deviations for the bare pore are significant, they appear to be systematic, pointing to a systematic error between TEM-imaged pores and experimental measurements. However, the difference between unmodified and chemically-modified pores is consistent, with a 1.6 nm mean decrease in diameter after coating, which supports our interpretation that a successful coating was achieved. **Figure 2c** shows current traces of two 7-nm SiN_*x*_ pores, one silanized (purple) and one bare (black). As expected, the silanized pore has less ionic conductance yet is stable and has larger low-frequency noise, as shown by the power spectral density (PSD) plots at 100 mV (inset). Finally, in accordance with previous studies,^45, 46^ a common outcome of chemical modification of a pore is that molecular fluctuations, formation of transient hydrophobic pockets, and charge fluctuations, all cause increases in 1/*f* noise.

As we have shown in our previous hybrid system,^36^ successful portal protein insertion and corking into a SiN_*x*_ nanopore with desired orientation required maintaining a voltage bias across the membrane to electrokinetically drive the protein to the SiN_*x*_ nanopore whose geometry is commensurate with the protein’s structure and size (**Figure 1d**). The charge bipolarity of the portal protein’s external surface^37^ prompts its orientation in the direction of the electric field, but the overall net force on the portal is such that reversing the bias results in ejection of the portal from the pore, limiting the applicability of our hybrid system at various applied voltages. In our current study, we have implemented the exact mechanism for the formation of the hybrid nanopore system along with chemically fixing the protein to the SiN_*x*_ nanopore to inhibit its uncorking in reverse voltage and attain permanent immobilization of the portal protein within the support membrane. As shown in **Figure 3a**, insertion of the protein into the silanized SiN_*x*_ nanopore is observed by a drop in the ionic current. We confirmed the successful disulfide bridge formation between the portal protein and the thiol groups on the pore walls by reversing the voltage bias to -400 mV, which typically did not result in ejection of the portal protein from the SiN_*x*_ nanopore. This indicates a stable hybrid nanopore with applicability in both voltage biases. An unsuccessful hybrid nanopore formation is easily identified as a sudden increase in the ionic current to an “open pore” conductance level when the voltage is reversed, as shown in **Figure 3b. Figure 3c** shows a current-voltage curve of a 7 nm silanized SiN_*x*_ pore before and after hybrid nanopore formation. As seen in the current-voltage curve, the hybrid nanopore is stable at high voltages (up to 500 mV shown here). The critical component in successful disulfide bridge formation between portal protein and silanized SiN_*x*_ pore is the use of the catalyst Cu(phenanthroline)_2_, which facilitates the thiol oxidation through a previously reported mechanism^47^ (shown in **Figure 3d**). We did not observe any successful disulfide hybrid pore disulfide bridge formation in the absence of this catalyst. Moreover, we speculate that the arbitrary geometry of the fabricated SiN_*x*_ pores and the deviation of their diameters, as shown in **Figure 2b**, render the geometry of some pores unfit for successful portal protein insertion or disulfide bridge formation.

**Figure 3.**
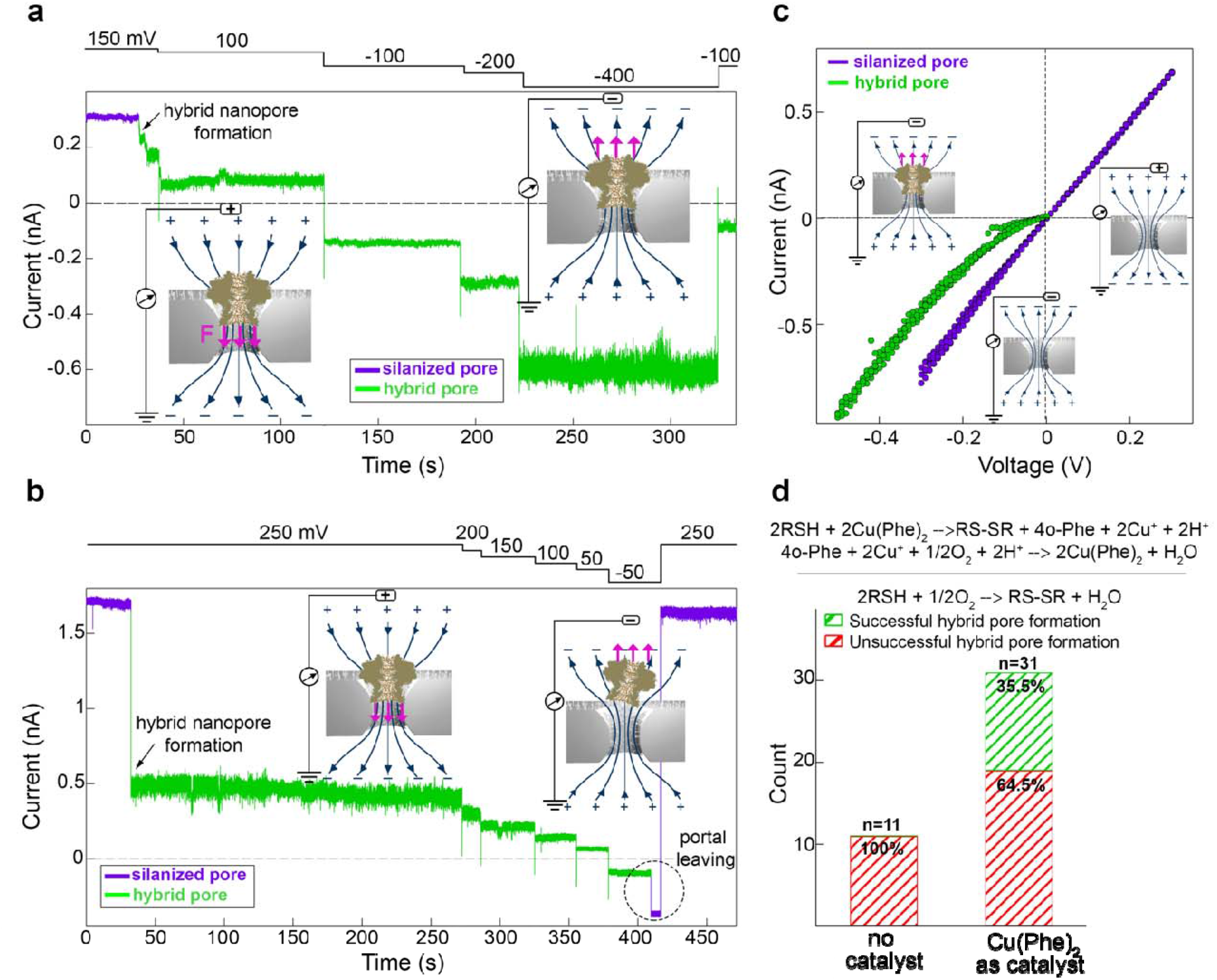
Stable hybrid nanopore formation and characterization. (a) An example of a successful hybrid nanopore formation through disulfide bridge formation between the portal protein and thiolated nanopore surface. The sudden drop in the ionic current indicates the protein insertion into the SiN_*x*_ pore which is also stable at the reverse voltage. The schematic shows the direction of the applied force on the protein in each electric field. Traces were recorded at 100 kHz sampling frequency and lowpass filtered at 10 kHz. Buffer: 0.5 M NaCl, 20 mM HEPES, pH 7.5, 160 mM Cu(phenanthroline)_2_ (b) An example of unsuccessful hybrid nanopore formation as indicated by ejection of the portal protein from the SiN_*x*_ nanopore by a sudden increase in the ionic current at the reverse voltage. Traces were recorded at 250 kHz sampling frequency and lowpass filtered at 10 kHz. Buffer: 0.5 M NaCl, 20 mM HEPES, pH 7.5, 320 mM Cu(phenanthroline)_2_ (c) Current-voltage curve of a 7-nm diameter silanized SiN_*x*_ nanopore, before and after portal protein insertion. This graph shows the stability of our hybrid system up to 500 mV. (d) Success rate for chemically bound hybrid nanopore formation with and without the use of a catalyst (catalytic reaction depicted above the plot). Unsuccessful hybrid pore formation is defined by portal ejection from the SiN_*x*_ pore upon voltage reversal, as shown in (b).

We tested our stable hybrid nanopore system’s capability for sensing biomolecules through studying ssDNA translocation in both crown-to-clip and clip-to-crown directions. In our previous study we only probed biomolecule translocation in the clip-to-crown direction due to the corked portal hybrid pore’s instability upon reversal of voltage polarity.^36^ **Figure 4a** shows a schematic of the experiment where negative voltage is applied to the *trans* chamber to facilitate ssDNA translocation in the crown-to-clip direction. **Figure 4b** shows 5-second current traces of the hybrid nanopore at -200 mV, -300 mV, and -400 mV, where transient drops seen in the ionic current indicate temporary occlusions of the nanopore by the ssDNA molecules. **Figure 4c** shows a close-up look at three representative events for each voltage. To confirm ssDNA molecules transport through the hybrid nanopore in the crown-to-clip direction, an analysis of the event characteristics is required. **Figure S3** shows scatter plots of ionic current blockage versus dwell time from -140 mV to -400 mV in 20 mV steps. The plots show that at voltages < 220 mV, the events are primarily low-level current blockades with spread-out dwell times indicating collision of DNA molecules with the portal protein and failed translocation. While not serving as concrete proof, this behavior is consistent with a recent structural study which indicated the portal may serve as a one-way valve preventing dsDNA leakage from capsids during virus assembly^48^. The mechanism involves “loop-in” conformation activated during transport in the crown-to-clip direction, when portal’s crown adjustement triggered by DNA forces the internal tunnel loops in the pore lumen to extend towards DNA and prevent its translocation along the tunnel. Our observed behavior in the low-voltage regime is also consistent and a former study on membrane-embedded portal with a similar portal mutant which showed that β-cyclodextrin transport does not occur in the crown-to-clip direction for the voltage range studied (<140 mV).^37^ In contrast to the low voltage behavior, at voltages higher than 220 mV the scatter plots reveal a new population of events with deeper blockades and dwell times that decrease with increasing voltage. To study the dwell time change as a function of voltage, we fitted exponential functions to the dwell time distribution shown in **Figure S4. Figure 4d** shows that from 140 to 220 mV, the mean dwell time of events increases with voltage, while from 220 to 400 mV, dwell time decreases with voltage. This suggests that below a certain voltage threshold (220 mV here) ssDNA molecules only collide with the portal protein, while above the threshold ssDNA molecules start to fully transport through the pore in the crown-to-clip direction. We note that at voltages above ∼ 320 mV, a new (minor) population of slow events emerges, which we attribute to ssDNA/protein interaction or sticking (not shown in **Figure 4d**, see **Figure S5**). We also studied the dynamics of ssDNA translocation in the clip-to-crown direction by applying a positive voltage to the *trans* chamber and adding DNA to the cis chamber (**Figure S6a). Figure S6b** show 5-second current traces of the hybrid nanopore after adding ssDNA at 40 mV, 60 mV, 80 mV, and 100 mV. Analysis of the events (**Figures S6c** and **S7a**) show that at all voltages, there is a population of fast events that does not depend on voltage which is characteristic of ssDNA collision with the portal protein. At 80 mV and 100 mV, the second population of events with higher dwell time emerges, which we attribute to DNA transport through the portal protein in the clip-to-crown direction. The results demonstrate that there is less threshold for ssDNA molecules to transport in the clip-to-crown direction than in the crown-to-clip direction. Analysis of the capture rate as a function of voltage (**Figure S7b**) depicts an increase in the rate with increasing voltage.

**Figure 4.**
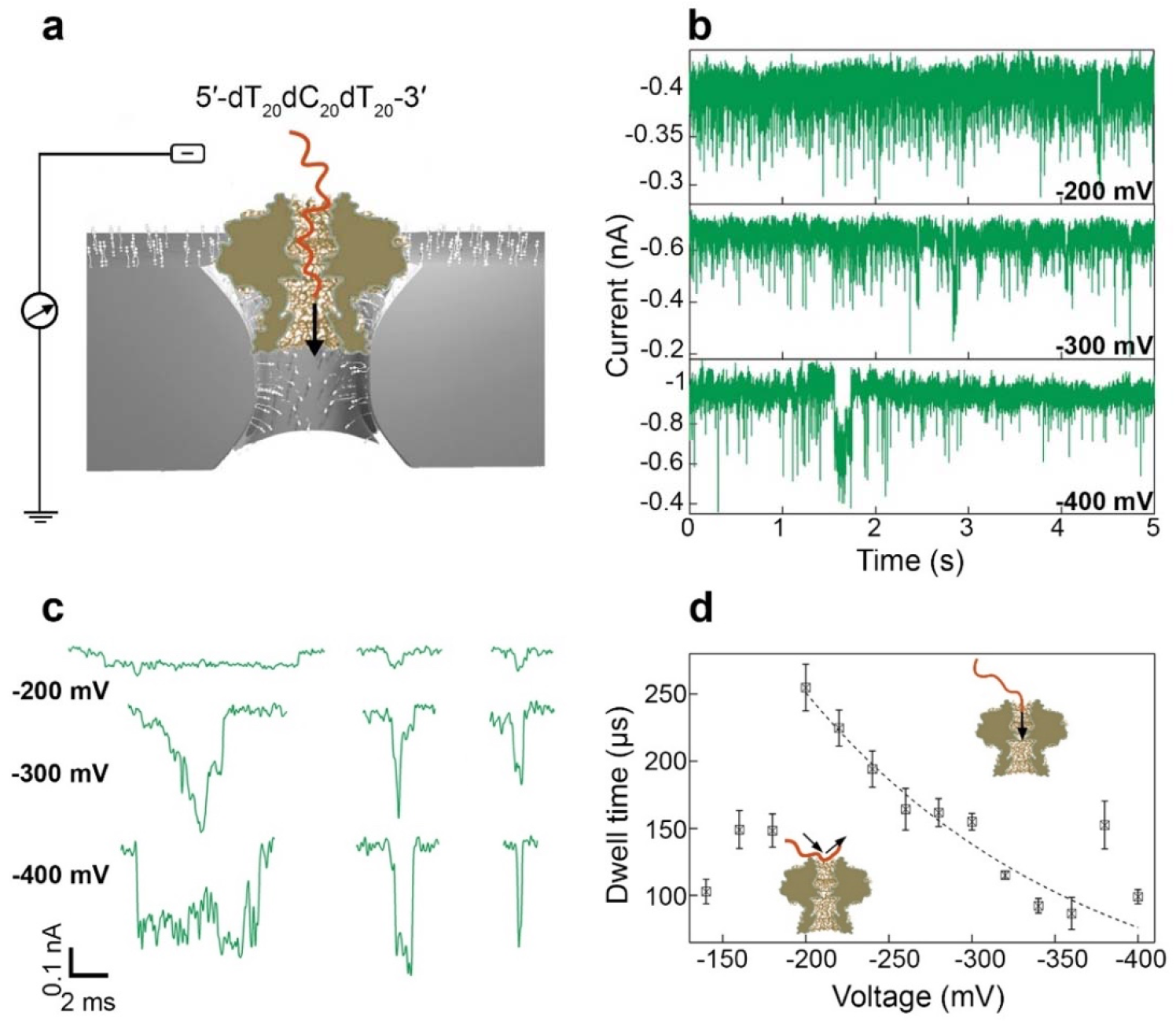
Single-stranded DNA transporting through the hybrid nanopore in the crown-to-clip direction. (a) Schematic of experimental setup. (b) 5-second current traces at -200 mV, -300 mV, and -400 mV demonstrate transient drops of the ionic current, characteristic signals of nanopore occlusion by DNA molecules (ssDNA final concentration: 6.25 μM). Traces were recorded at 100 kHz sampling frequency and lowpass filtered at 10 kHz. (c) Three representative events at -200 mV, -300 mV, and -400 mV. (d) Mean dwell time as a function of voltage showing that below -200 mV, ssDNA molecules only collide with the portal protein while at voltages higher than 200 mV DNA ssDNA molecules transport through the portal protein in the crown-to-clip direction as confirmed by the decrease in the dwell time as the applied voltage increases. Dwell time average values were calculated from the exponential fit to the distributions (**Figure S5**). The dashed line shows an exponential fit to the points. Data point at 380 mV is an outlier and was excluded in the fitting process. Buffer: 0.5 M NaCl, 20 mM HEPES, pH 7.5, 160 μM Cu(phenanthroline)_2_.

To further test our hybrid system for high-voltage sensing and sequencing applications, we studied DNA ratcheting through the hybrid nanopore using an Oxford Nanopore Technologies sequencing chemistry. **Figure 5a** shows a schematic of the experiment where a motor protein bound to its DNA molecule rests on the portal protein and ratchets its DNA through the pore. We carried out the experiment at voltages from -200 mV to -750 mV to demonstrate the activity of our hybrid system at high voltage. **Figure 5b-f** show current traces from -200 mV to -600 mV. On the right side, a close-up view of the highlighted regions is shown. At 200 mV, distinct features correlated with the stepwise motion of the DNA through the portal protein are visible, while at higher voltages these features become less frequent (**Figure S8** shows more examples of events). **Figure 5g** demonstrates scatter plots of current blockage *vs*. dwell time at 300 mV and 400 mV for both freely transporting ssDNA and motor protein-bound DNA molecules. This side-by-side comparison shows that the motor protein feeds DNA into the pore ∼ 5 orders of magnitude slower. Moreover, as indicated by current traces at different voltages and dwell time decrease as a function of voltage, DNA strips off from the motor protein at high voltages (> 450 mV) due to the high electrostatic force intervening with the ratcheting process (**Figure S9**). Therefore, although our hybrid nanopore system shows stable activity up to 750 mV, the motor protein is not optimized for use at high voltages. In a control experiment using motor protein/DNA with silanized SiN_*x*_ nanopore (**Figure S9**) at high voltage (> 600 mV), signals are less frequent and completely distinct from equivalent signals in the hybrid nanopore, which confirms that these signals are indeed due to DNA ratcheting through the portal protein.

**Figure 5.**
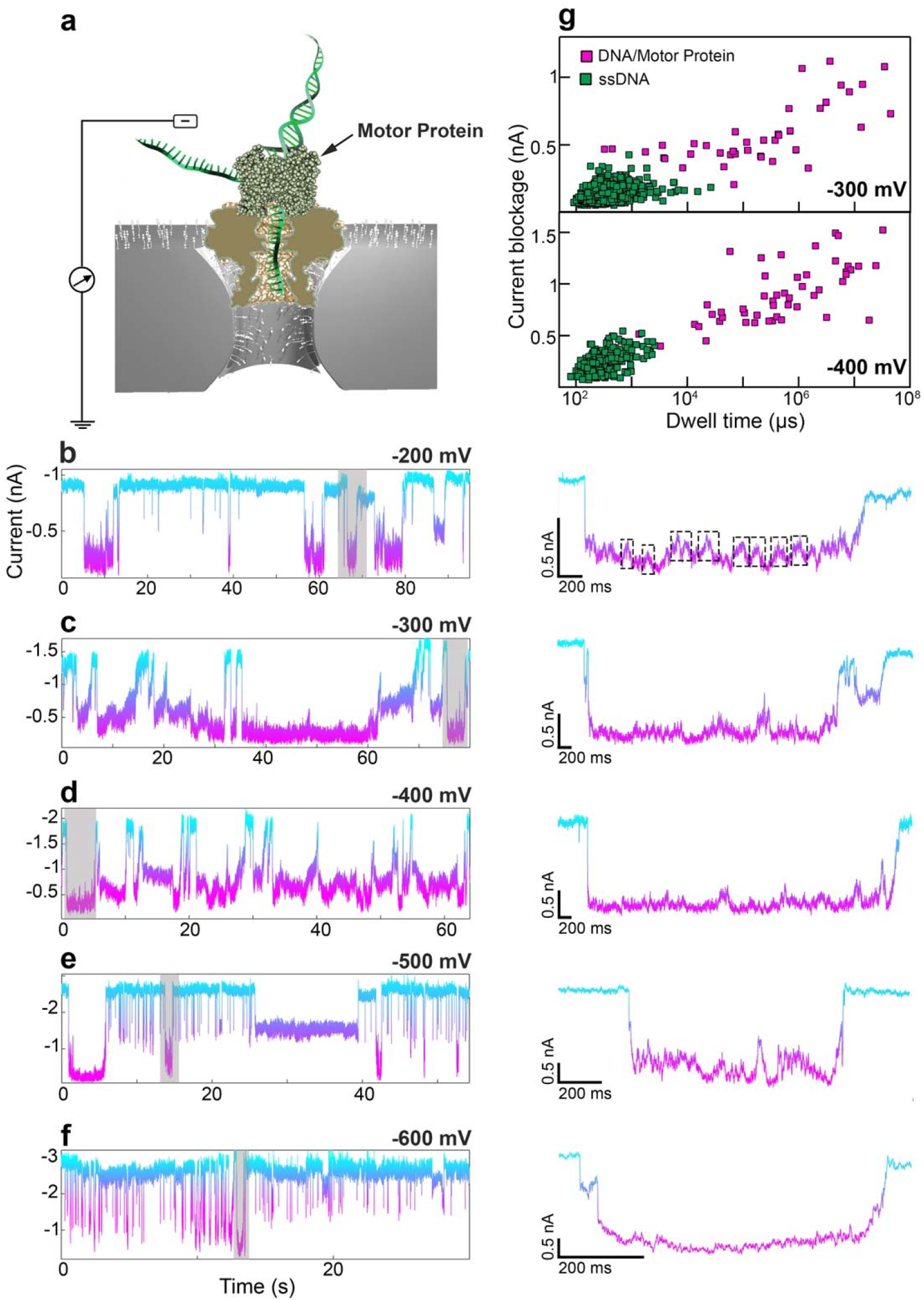
Single-stranded DNA ratcheting through the hybrid nanopore in the crown-to-clip direction. a) Schematic of our experimental set-up for motor protein-controlled DNA transport through a hybrid pore. (b) Current traces at -200 mV, -300 mV, -400 mV, -500 mV, and -600 mV, demonstrating transient drops of the ionic current, which we attribute to DNA molecules ratcheting through the portal protein (traces were digitized using a sampling rate of 100 kHz after lowpass filtering at 10 kHz). To the right of each trace is a close-up view of the highlighted regions at each voltage. The traces were further lowpass filtered at 2 kHz. At 200 mV, the dashed lines highlight the stepwise change in the ionic current, which we attribute to DNA ratcheting motion. (g) Scatter plots of current blockage versus dwell time at 300 mV and 400 mV to demonstrate a side-by-side comparison of DNA transport dynamics with and without ratcheting. Buffer: 0.5 M NaCl, 20 mM HEPES, pH 7.5, 320 μM Cu(phenanthroline)_2_, 75 mM KCl, 7 mM ATP, 7 mM MgCl_2_.

## Conclusion

In summary, we demonstrated that by chemically fixing the *G20c* portal protein within a synthetic support membrane, a highly robust hybrid nanopore platform with voltage stability of up to 750 mV could be attained. Moreover, we showed high-voltage recognition of freely translocating and motor protein-mediated DNA ratcheting through the hybrid nanopore, with the motor protein slowing down the DNA transport down to five orders of magnitude than the electrophoretically-driven DNA transport. Developing this high-voltage system further could significantly improve current nanopore sequencing platforms by providing superior resolution due to the higher potential signal-to-noise values in this hybrid pore system; the ability to apply higher voltage then in conventional organic membranes also offers opportunities to re-read molecules by retracting them electrically and then re-engaging the enzyme motor. Future work will focus on reducing the noise properties of this hybrid system, as well as assessing the portal protein’s sensing region and its resolution limitations for biopolymer sequencing.

## Materials and Methods

### Protein Cloning, Expression, and Purification

CD/N mutant portal protein was expressed and purified as explained previously.^36^ In summary, this mutant was expressed in Escherichia coli shuffle cells after reaching OD600 = 0.8 and induction with 0.5 mM isopropyl β-D-1-thiogalactopyranoside (IPTG) at 30 °C overnight. Afterward, the cell pellets were resuspended in 1 M NaCl, 50 mM Tris pH 8, 10 mM imidazole, 2 mM DTT, and kept at -80 °C until purification. For purification, the cells were thawed, lysed by sonication after addition of 10 mg.ml^−1^ lysozyme, protease inhibitor tablets (Thermo Scientific, A32963), and further clarified by centrifugation at 12k rpm for 55 minutes. Protein purification included 6 steps: 1. purification by Immobilized Metal Affinity Chromatography (IMAC; 5 mL HiTrap FF Crude, GE Healthcare), 2. buffer exchange to 0.5 M NaCl, 50 mM Tris pH 8, 50 mM potassium Glutamate, 1 mM DTT using a desalting column (HiPrep 26/10; GE Healthcare), 3. overnight cleavage of the histidine affinity tag using 3C protease, 4. buffer exchange into 1 M NaCl, 50 mM Tris pH 8, 10 mM Imidazole, 2 mM DTT using the same desalting column, 5. purification by IMAC for the second time to remove histidine-tagged proteins, 6. purification by Size Exclusion Chromatography (SEC; Superose 18/300; GE Healthcare). The purified proteins were flash-frozen and kept at -80 °C until use.

### Silicon Nitride Membrane Fabrication

50-nm-thick free-standing SiN_*x*_ membranes were fabricated at the center of 5 mm × 5 mm silicon chips. In summary, 50-nm-thick, high-stress (250 MPa) SiN_*x*_ film was deposited on a 2 µm-thick SiO_2_ layer/300 µm-thick silicon wafers. Photolithography (Karl Suss MA6 mask aligner) and several wet and dry etching steps were used to pattern the final 20 µm × 20 µm SiN_*x*_ membranes. 3 µm circular areas on the membrane (known as thin regions) were locally thinned to 25-30 nm using Technics Micro-RIE Series 800 for SF_6_ plasma etching (200 mTorr, 50 W, 40 s) and further confirmed by AFM.

### Solid-State Nanopore Chemical Modification

7-8 nm nanopores were fabricated through the thin region of the SiN_*x*_ membranes using TEM. Each chip containing the membrane and fabricated pore was thoroughly cleaned first by immersing in hot piranha solution (1:2 H_2_O_2_:H_2_SO_4_) for 30 minutes and hot water for 30 minutes for any organic residue and contamination to be removed and the surface to be rendered hydrophilic and hydroxylated (piranha solution was always used freshly prepared in glassware while kept inside a fume hood). Afterward, chips were dried entirely using an N_2_ gun and baked at 150 ºC for 3 minutes. The silanization protocol was adapted from the previous work by Kim *et al*.^42^ Each chip was placed into a 1.5 ml Eppendorf vial and moved into the glovebox with running N_2_ gas. 400 µl extra dry dichloromethane (Acros Organics) and 100 µl 2,2-dimethoxy-1-thia-2-silacyclopentane (Gelest, Inc.) were added to each vial and left for the silanization to complete for 1 - 3 hours. To remove the chemisorbed molecules on the surface, each chip was immersed in 5 vials of DCM and gently agitated. Each chip was kept in anhydrous ethanol (Acros Organics) until use and was dried before loading on the fluidic cell. Silicone elastomer was used to seal the chip edges. Ethanol was used to facilitate wetting of the silanized pores.

### Nanopore Experiment Data Acquisition and Analysis

Axopatch 200B amplifier at either 100 kHz or 250 kHz sampling rate was used for single-channel measurements. After wetting the silanized pores using ethanol and getting a stable current, portal protein with a final concentration of ∼ 20 μg/μl was added to the trans chamber and Cu(phenanthroline)_2_, with a final concentration of 160 μM - 320 μM, was added to both chambers. After portal protein insertion into the SiN_*x*_ nanopores, detected by a sudden drop in the ionic current, voltage was reversed to check the protein’s stability. If the protein was stable at both voltages and the hybrid nanopore was successfully formed, the desired analyte was added to the trans or cis chamber. Oxford Nanopore rapid sequencing kit (SQK-RAD004) was used for motor protein/DNA experiments. Data processing was done using OpenNanopore software (https://lben.epfl.ch/page-79460-en.html) and Pythion software (https://github.com/rhenley/Pyth-Ion/) and further analyzed using Igor Pro software. For dwell time distributions, the first two bins representing events < 150 µs were excluded in the fitting process due to the underrepresentation of events close to the 10 kHz filtering threshold.

## Notes

The authors declare no competing financial interest.

## Acknowledgment

The authors would like to acknowledge Oxford Nanopore Technologies plc, Oxford, United Kingdom for financial support of this work, as well as early-stage support from the Bilateral United Kingdom Biotechnology and Biological Sciences Research Council (BBSRC) – United States National Science Foundation (NSF) Lead Agency Pilot Program grant (BB/N018729/1 and NSF-1645671 to A.A.A. and M.W., respectively). The authors also acknowledge Dr. Min Chen for constructive discussions.

