## Supplementary material for "High-Voltage Biomolecular Sensing Using a Bacteriophage Portal Protein Covalently Immobilized Within a Solid-State Nanopore": SI

**Table of Contents**


**Figure S1.** 2D class averages calculated using cryo-EM images of CD/N mutant of the *G20c* portal protein. (page 3)

**Figure S2.** Functionalization of $SiN_x$ surface with MPTES (page 4).

**Figure S3.** Scatter plots of current blockage versus dwell time for events from -140 mV to -400 mV in 20 mV steps (page 5).

**Figure S4.** Dwell time histogram of events from -140 mV to -400 mV in 20 mV steps (page 6).

**Figure S5.** Mean dwell time as a function of voltage for the second population of events shown in Figure S3 (page 7).

**Figure S6.** ssDNA transport through the hybrid nanopore in the clip-to-crown direction (page 8).

**Figure S7.** Analysis of ssDNA transport dynamics in the clip-to-crown direction (page 9).

**Figure S8.** Examples of individual events for DNA feeding through the portal protein by the motor protein (page 10).

**Figure S9.** Scatter plots of current blockage versus dwell time for events from -200 mV to -750 mV in 50 mV steps (page 11).

**Figure S10.** ssDNA ratcheting through the chemically modified $SiN_x$ nanopore (page 12).

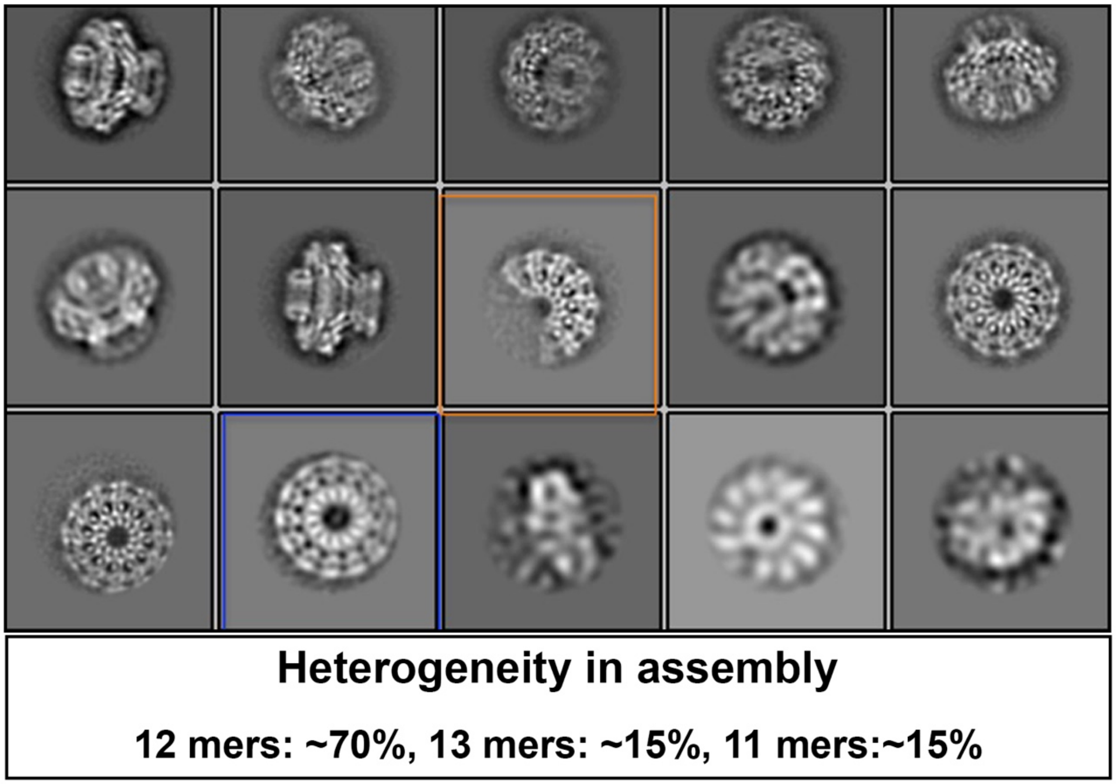


**Figure S1. 2D class averages calculated using cryo-EM images of CD/N mutant of the**

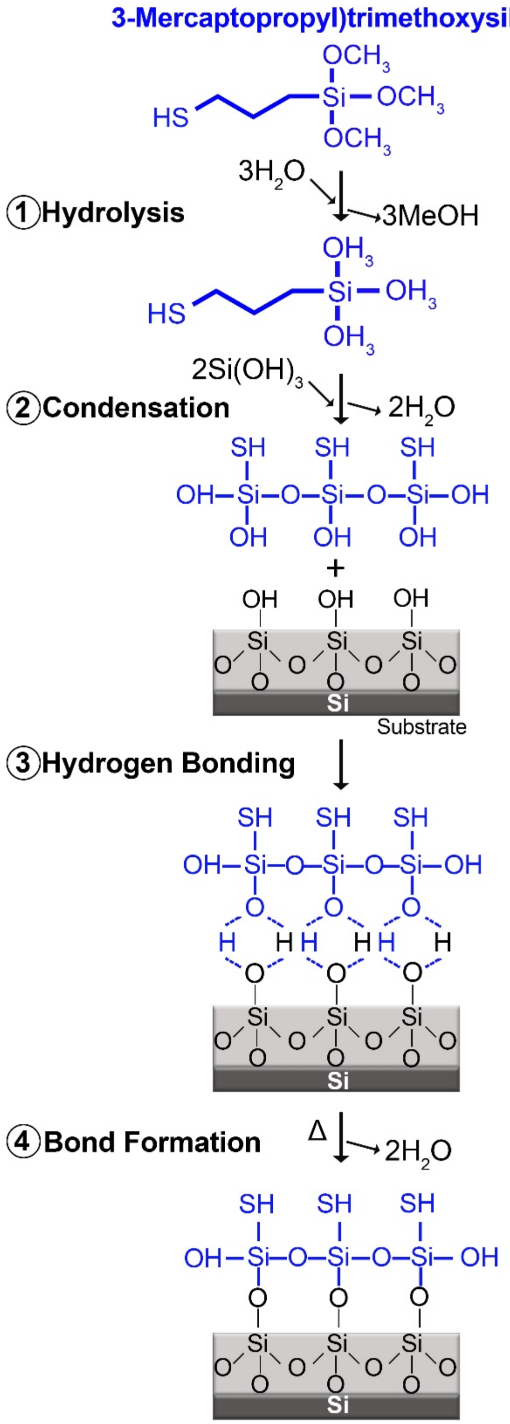


**Figure S2. Functionalization of SiN$_x$ surface with MPTES.**

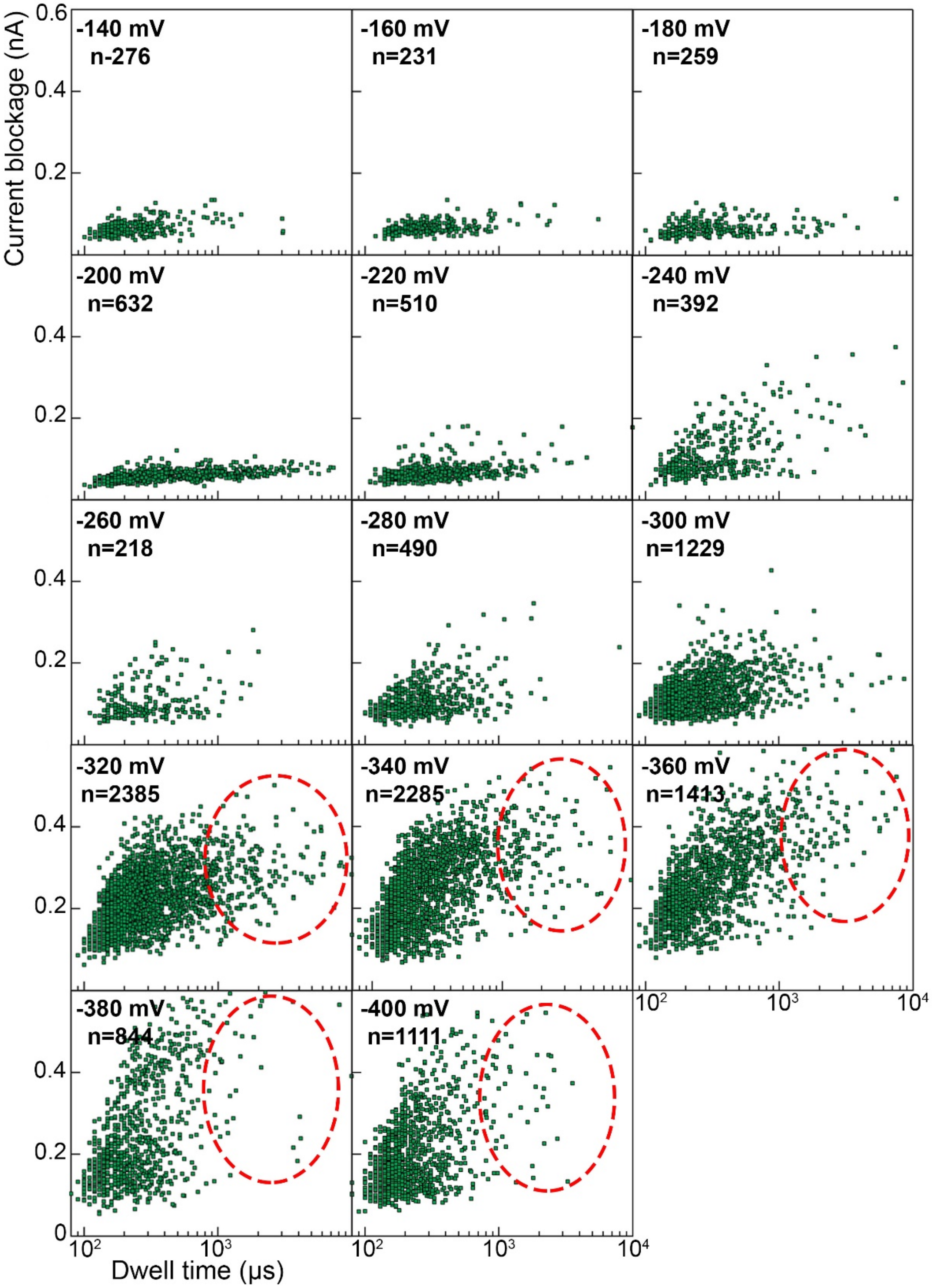


**Figure S3. Scatter plots of current blockage versus dwell time for events from -140 mV to -400 mV in 20 mV steps.** At voltages higher than -300 mV a second population of events emerge shown

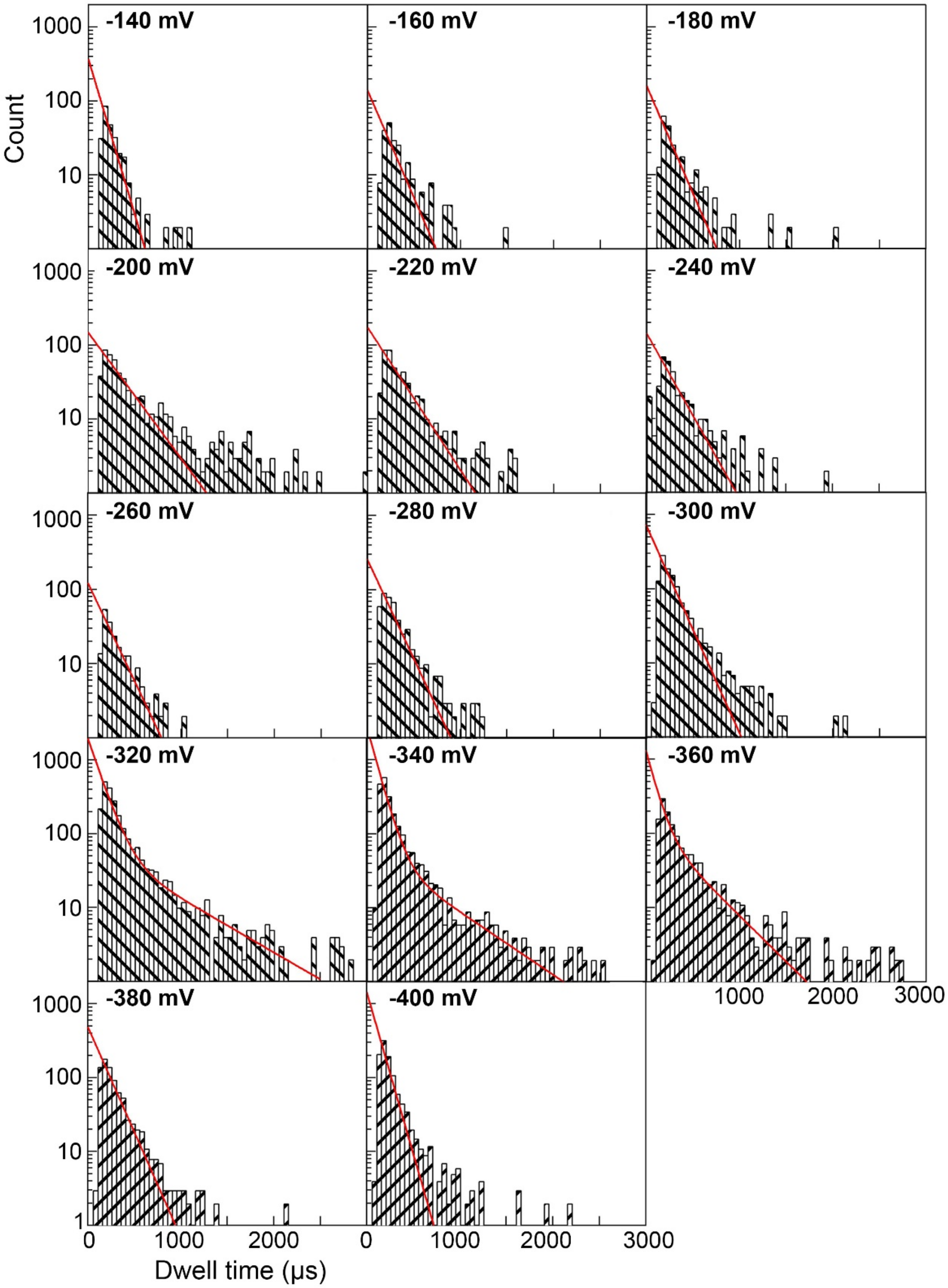


**Figure S4. Dwell time histogram of events from -140 mV to -400 mV in 20 mV steps.** Red lines show exponential fits to the histograms. Starting from -320 mV, as the second population of events start to emerge, we fitted double exponential function. However, there are not enough data points at -380 mV

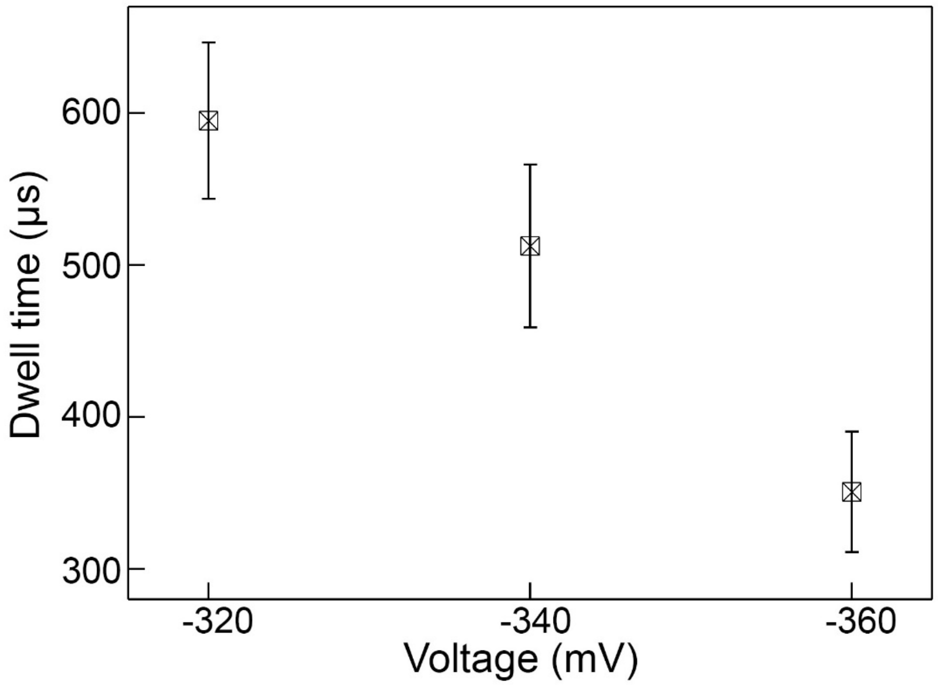


**Figure S5. Mean dwell time as a function of voltage for second population of events shown in Figure S4.** For -380 mV and -400 mV there were not enough data points to fit the distribution to the

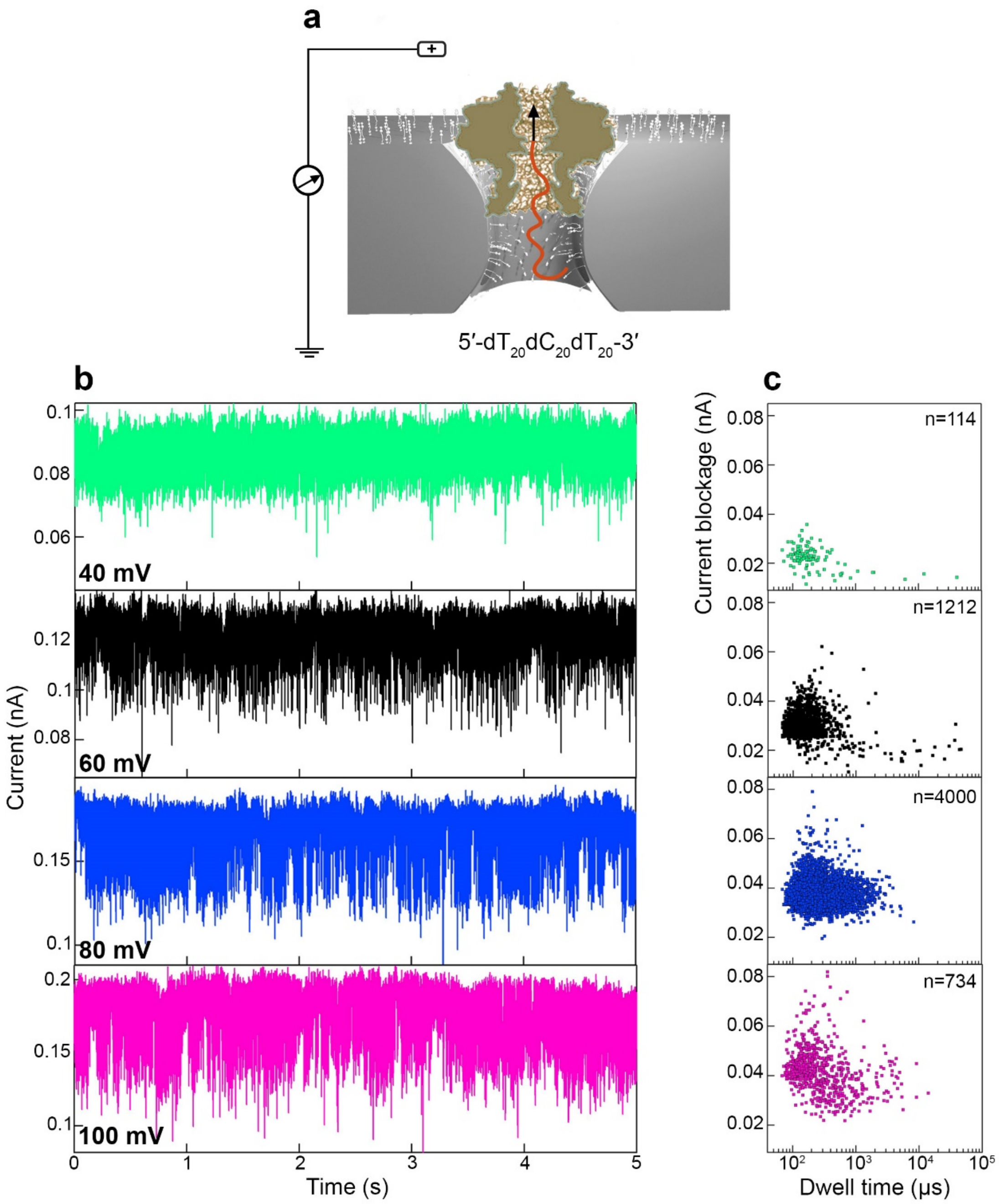


**Figure S6. ssDNA transport through the hybrid nanopore in the clip-to-crown direction.** (a) Schematic of our experimental set-up. (b) 5-second current traces at 40 mV, 60 mV, 80 mV and 100 mV, demonstrating transient drops of the ionic current, characteristic signals of nanopore occlusion by DNA molecules (ssDNA concentration: 10 μM). Traces were recorded at 250 kHz sampling frequency and low-pass filtered at 10 kHz. (c) Scatter plots of current blockage versus dwell time for events from M

$Cu(Phenanthroline)_2$.

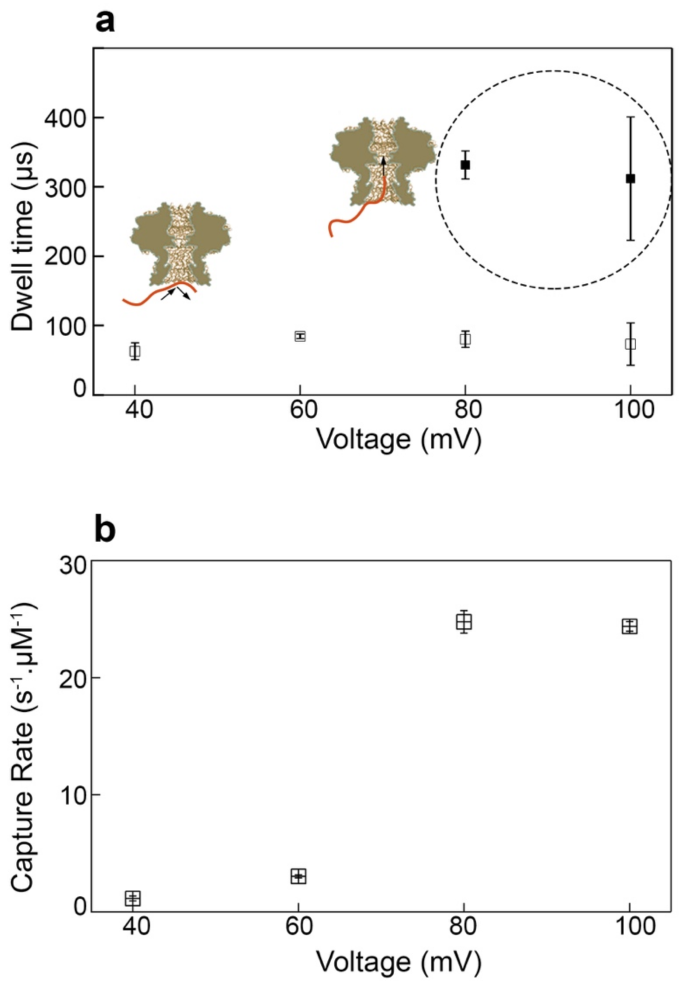


**Figure S7. Analysis of ssDNA transport dynamics in the clip-to-crown direction.** (a) Mean dwell time as a function of voltage showing transport of DNA molecules through the portal protein occurs at higher voltages than 80 mV. (b) Normalized capture rate as a function of voltage shows increase in the capture rate by increasing voltage. Average values were calculated from the exponential fit to the

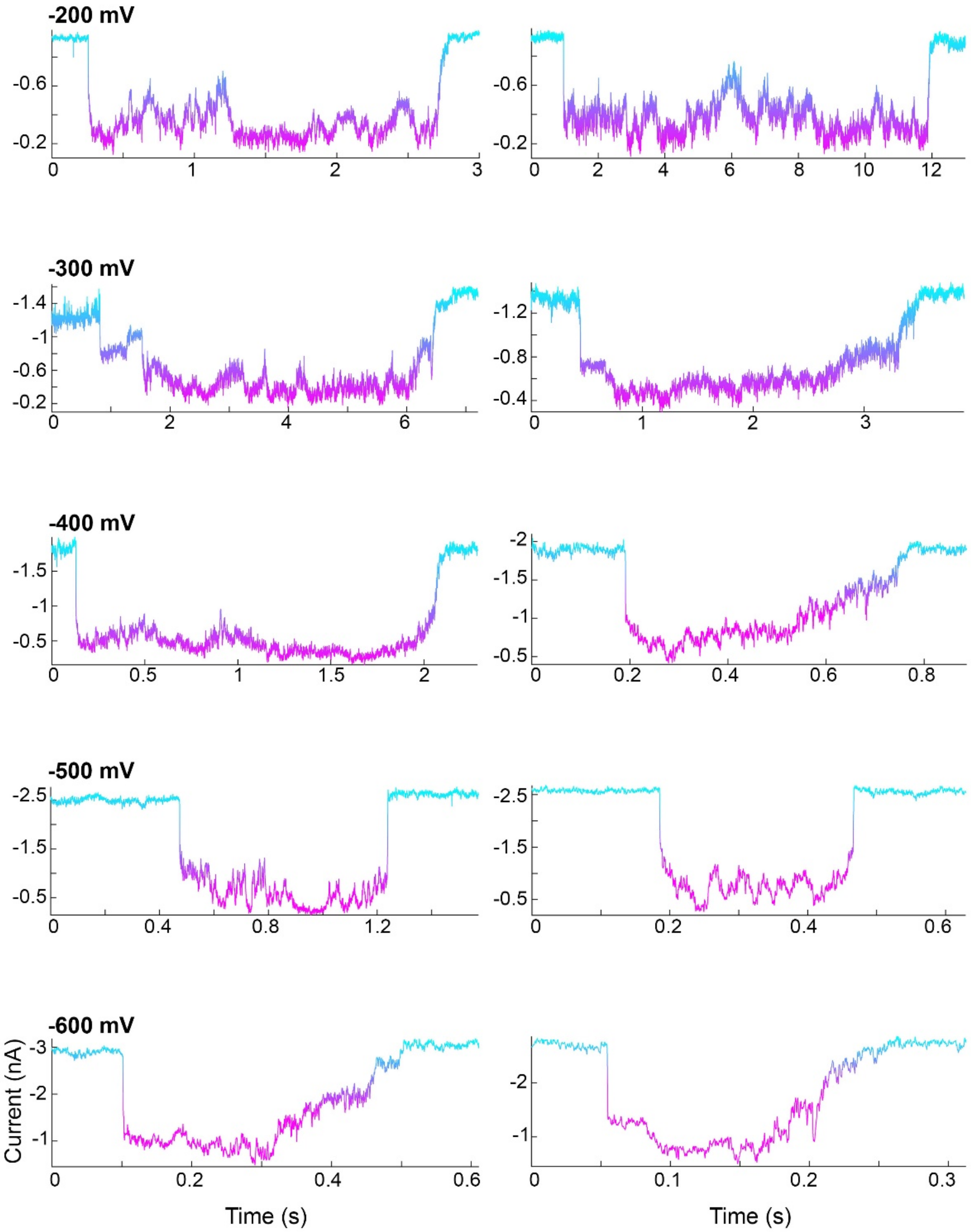


**Figure S8. Examples of individual events for DNA feeding through the portal protein by the motor**

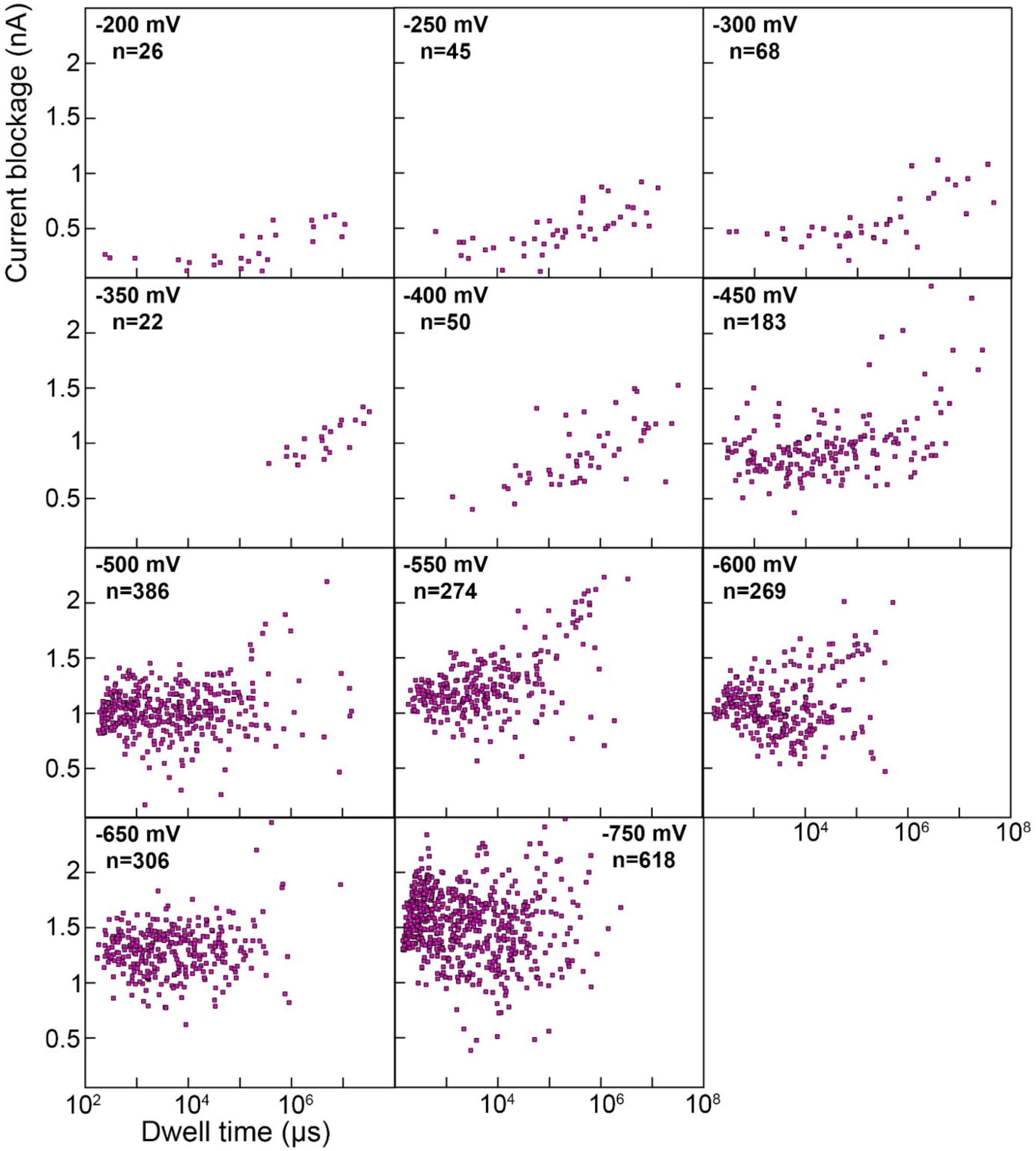


**Figure S9. Scatter plots of current blockage versus dwell time for events from -200 mV to -750**

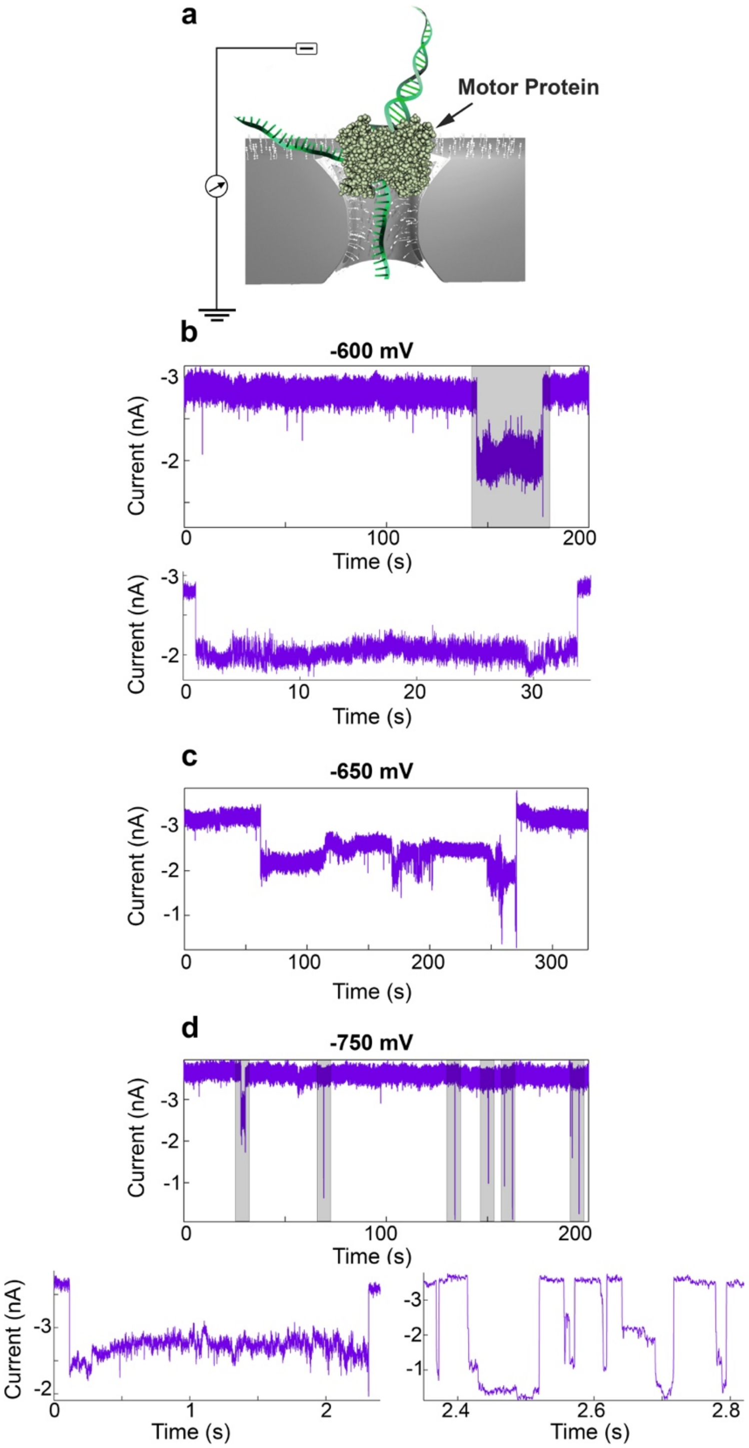


**Figure S10. ssDNA ratcheting through the chemically modified $SiN_x$ nanopore.** a) Schematic of our experimental set-up. (b) Current trace at -600 mV. The bottom trace shows the zoomed-in view of the highlighted events. (c) Current trace at -650 mV. (d) Current trace at -750 mV. The bottom traces show the close-up look of the highlighted regions. Traces were recorded at 100 kHz sampling frequency. Main traces were low-pass filtered at 10 kHz and the events were further filtered at 2 kHz. Buffer: 275
